# Is hidden demographic heterogeneity pervasive? Harvested birds as an example

**DOI:** 10.64898/2026.11.09.764468

**Authors:** Emilienne Grzegorczyk, Camille Schatz, Charlotte Francesiaz, Roger Pradel, Christophe Barbraud, Guillaume Souchay, Alain Caizergues, Kévin Le Rest, Matthieu Guillemain, Jocelyn Champagnon, Cyril Eraud

## Abstract

Individual demographic heterogeneity refers to the fact that individuals from a given population exposed to similar environmental conditions express different demographic traits or tactics. This heterogeneity can either be linked to factors/states such as age, sex or body condition, or be due to hidden factors/states not immediately apparent or easily characterized (hereinafter referred to as ’hidden heterogeneity’). Yet accounting for demographic heterogeneity, and more particularly heterogeneity in survival is crucial because it is an important mechanism underlying compensation processes between natural and harvest mortality. In this study, we used capture-mark-recapture-recovery data of 6 hunted species covering a large range of the demographic slow-fast continuum (61,432 individuals ringed from 1953 to 2020 among Common Teal *Anas crecca*, Mallard *A. platyrhynchos*, Common Pochard *Aythya ferina*, Common Snipe *Gallinago gallinago*, Common quail *Coturnix coturnix,* and Blackbird *Turdus merula*), to assess the presence of unexplained heterogeneity in survival with multievent models. We detected hidden survival heterogeneity in most species, even though the differences between the high and low categories of survival were often small. The probability for an individual to display a low vs high survival rate was not related to potentially important individual factors/states such as migration phenology or to morphology. The presence of hidden heterogeneity was pervasive, affecting species distributed along a relatively large spectrum of demographic strategies. This, in our opinion, supports that part of this heterogeneity in survival evidenced in our study is indeed hidden, resulting either from hidden intrinsic (inherent) individual differences of survival among individuals, or hidden extrinsic factors e.g., linked to spatial variations in hunting pressure. Our findings suggest that hidden heterogeneity needs to be explored more thoroughly in huntable species and accounted for in management policies.

## INTRODUCTION

Natural selection is the evolution engine whose fuel is the heritable heterogeneity between individual (Darwin, 1859). Individuals within populations usually differ in many respects including morphology (e.g., Grzegorczyk et al., 2022), physiology (Cohen & Hamrick, 2003, Biro & Stamps, 2010) and behavior (Shelton & Martins, 2017, McMahon et al., 2022). Part of these differences are genetically inherited and correlated with life-history or demographic traits or strategies and hence affect fitness (e.g., Koehl, 1996, Réale et al., 2007, Réale et al., 2010, Kern et al., 2016, Goulet et al., 2018, Gaona-Gordillo et al., 2023). Characterizing and quantifying such individual differences in demographic traits and strategies is a key step towards a better understanding of how natural populations resist, adapt or succumb to anthropogenic or natural stressors, and hence a prerequisite to any conservation actions (e.g., Stearns, 1976, Williams, 2013, Koons et al., 2014). Demographic strategies range along a continuum, from fast species that rely primarily on reproduction but are short-lived, to slow species that rely on high survival (are long-lived) but produce fewer offspring annually (Stearns, 1983, Lebreton & Clobert, 1991, Koons et al., 2014). Species position along this continuum allows one to evaluate their overall sensitivity to environmental changes such as the consequences of human activities including climate change (Wright & Trippel, 2009, Servanty et al., 2011, Stanton et al., 2018, Dunn & Møller, 2019), depending on whether such activities are affecting survival versus fecundity (Williams, 1996). Assessing demographic strategies can also be used to evaluate the extinction risk of a population facing threats (McLaughlin et al., 2002, Yarra & Magoulick, 2019) or to build scenarios by projecting population trends depending on the management decisions (either to enhance survival or reproduction e.g., Bakker et al., 2009, Wolf et al., 2010).

Much effort has been devoted to the study of life history/demographic tactics (Caswell & Vindenes, 2018, Gimenez et al., 2018, Hamel et al., 2018). Many of these studies have focused on survival, often showing some degree of heterogeneity among populations within species (e.g., Folliot et al., 2020) and, between individuals within populations (Guillemain et al., 2014, Gimenez et al., 2018, Folliot et al., 2020, Gibson et al., 2025). It results that much of the individual heterogeneity in survival can be explained either by intrinsic factors such as age, sex, social breeding, or migratory status (e.g., Lebreton et al., 1992, Weimerskirch, 1992, Martin, 1995, Jeugd & Larsson, 1998, Clutton-Brock & Isvaran, 2007, Toïgo et al., 2007, Arnold et al., 2012, Naef-Daenzer & Grüebler, 2016, Zúñiga et al., 2017, Folliot et al., 2020), or by extrinsic factors including site quality, weather conditions or other kinds of environmental irregularities (e.g., Eraud et al., 2009, Duriez et al., 2012). Because of the difficulty to predict or assess factors/states that would affect individual demographic traits/tactics, demographic models have mainly focused of the effects of age or sex, so far. Hence, part of the individual heterogeneity has resisted all attempts of explicit characterization, and, as a result, has been qualified as unexplained or hidden (Guillemain et al., 2014, Cam et al., 2016, Gimenez et al., 2018). Yet, individual heterogeneity directly affects fitness and fitness strategies (see above and Stearns & Kawecki, 1994), as well as the demographic properties of natural populations e.g. extinction risk (Kendall & Fox, 2002, Robert et al., 2003), demographic stochasticity (Vindenes et al., 2008, Caswell & Vindenes, 2018), and long-term population growth rate (Kendall et al., 2011, Vindenes & Langangen, 2015). Considering heterogeneity, even when hidden from us, is therefore essential to correctly assess evolutionary or demographic processes and to infer proper management decisions.

Four main sources of unexplained individual demographic heterogeneity are currently recognized in animals. Firstly, the notion of frailty state that ontogenic and genetic factors underly the co-occurrence, in populations, of poor- and high-quality individuals that display large differences in terms of demographic contribution to the next generation (e.g., Cooch et al., 2014, Cam et al., 2016, Gimenez et al., 2018). Secondly, the theory of pace-or-life (POL) syndromes, hypothesizes that part of the heterogeneity within populations would result from the presence of individuals achieving the same fitness over the long-term albeit through different (genetically inherited) paths i.e., demographic tactics along the slow-fast continuum (Réale et al., 2010). Moreover, these inherited demographic tactics would covary with other measurable -but most often not accounted for-traits, like physiology, behavior, to combination of traits such as those underlying personality (Réale et al., 2010, Santicchia et al., 2018). Thirdly, another part of the hidden individual demographic heterogeneity may result from environmental stochasticity, more particularly from interactions between the individuals and their environment, that are particularly difficult to characterize and quantify (e.g., spatial variations in predation/hunting pressure Vindenes et al., 2008). Finally, hidden individual demographic heterogeneity could be generated by chance alone, that is by stochastic demographic processes (neutral model of heterogeneity, Cam et al., 2016). It should be noted that the resistance of some of these sources/causes of demographic heterogeneity to formal characterization attempts may be provisional thanks to statistical and theorical (Réale et al., 2010) advances such as multievent capture-recapture models and POL syndrome theory (Pradel, 2005, Gourlay-Larour et al., 2014, Guillemain et al., 2014).

In the wake of previous studies (Aubry et al., 2011, Lindberg et al., 2013, Guillemain et al., 2014, Péron et al., 2016, Guéry et al., 2017, Arnold, 2021), we addressed the issue of hidden survival heterogeneity in hunted birds with contrasted ecology, migration strategies, demographic paces, and levels of exploitation (Hirschfeld et al., 2019). Our study models included two fast species displaying generation times shorter than 2 years, Common Quail (*Coturnix coturnix*, hereafter Quail) and Blackbird (*Turdus merula*), two intermediate species with generation times between to 2 and 3 years, Common Teal (*Anas crecca*, hereafter Teal) and Mallard (*A. platyrhynchos*), and two slower species with a generation time over than 3 years, Common Pochard (*Aythya ferina*, hereafter Pochard) and Common Snipe (*Gallinago gallinago*, hereafter Snipe). Previous studies of hidden survival heterogeneity have mostly focused on protected species, yet assessing this type of heterogeneity, whose compensation processes depend on individual heterogeneity (Lindberg et al., 2013, Cooch et al., 2014, Caudill et al., 2017, Arnold, 2021, Grzegorczyk et al., 2024), is of special relevance in harvested species.

Using multievent models, we first assessed the presence of hidden individual heterogeneity in survival focusing on adults, for the sake of simplicity, we then explored potentially causal factors, such as migratory strategy (migration phenology) and individual quality (body mass) by comparing the distribution of these variables among the high vs low survival group. Finally, we assessed if our results were compatible with the demographic buffering hypothesis, which states that, in order to maximize fitness, individuals should strive at buffering the demographic parameter to which population growth rate, and hence fitness, is the most sensitive (i.e., reproductive output for fast species and adult survival for fast species Koons et al., 2014, Hilde et al., 2020), as protected species do (Pfister, 1998, Péron et al., 2016).

## Methods

### Datasets

The general dataset for Teal, Mallard, Pochard, Snipe, Quail and Blackbird covered the period 1953-2020 (with differences among species) and included information capture, live and dead encounter histories for a total of 61,134 individuals (Table 1). Trapping methods differed among species: mainly mist-netting for Blackbird, Snipe and Quail and mainly baited traps for the duck species. All birds were ringed using a unique metal and numbered ring and their age and sex were determined by the ringers. All Pochards were also marked using a nasal saddle bearing an alpha-numerical code (Gourlay-Larour et al., 2013). For Teal and Mallard, ringing wasinterrupted at our study sites in the mid-1970s and only reinstated in 2002. Accordingly, the data for these two species were split into two datasets: one“historical” dataset (before early 2000s) and a “modern” dataset (after early 2000s; Table 1). For species other than ducks, ringing sites were distributed across the whole mainland French territories. For Teal and Mallard, the only ringing area was in the Camargue Wetlands, i.e. the Rhône delta in Southern France. For Pochard the ringing area was the Lake of Grand-Lieu close to the mouth of the Loire River, in Western France (Table 1). Encounters consisted in live recaptures (visual sightings anywhere and local physical recaptures for Pochard, and local physical recaptures only for the other species) as well as dead recoveries anywhere.

**Table 1 -.** Description of the various data sets for each species.

| Species and Period | Sex | Nb of ringed individuals | Site | Period | Ringing season |
|---|---|---|---|---|---|
| **Historical Teal** | Female | 6,330 | Camargue | 1953/1954 to 1977/1978 | 1<sup>st</sup> August year n – 31 March year n+1 |
|  | Male | 11,806 |  |  |  |
| **Modern Teal** | Female | 218 | Camargue | 2002/2003 to 2017/2018 | 1<sup>st</sup> Aug year n – 31 Mar year n+1 |
|  | Male | 375 |  |  |  |
| **Historical Mallard** | Female | 1,031 | Camargue | 1953/1954 to 1976/1977 | 1<sup>st</sup> Aug year n – 31 Mar year n+1 |
|  | Male | 2,154 |  |  |  |
| **Modern Mallard** | Female | 243 | Camargue | 2002/2003 to 2010/2011 | 1<sup>st</sup> Aug year n – 31 Mar year n+1 |
|  | Male | 647 |  |  |  |
| **Pochard** | Female | 328 | Lake of Grand-lieu | 2004/2005 to 2019/2020 | 1<sup>st</sup> Oct year n – 31 Jan year n+1 |
|  | Male | 524 |  |  |  |
| **Snipe** | Female | 3,000 | All France | 1998/1999 to 2019/2020 | 1<sup>st</sup> Sept year n – 30 Apr year n+1 |
|  | Male | 2,515 |  |  |  |
| **Quail** | Male | 12,095 | All France | 2009/2010 to 2019/2020 | 1<sup>st</sup> Apr year n - 15 Jul year n |
| **Blackbird** | Female | 10,424 | All France | 1988/1989 to 2017/2018 | 1<sup>st</sup> Oct year n – 31 Mar year n+1 |
|  | Male | 9,444 |  |  |  |

### Data formatting

For the purposes of the present analyses, only adults whose sex was recorded upon ringing were included, so as to simplify the analyses by limiting the sources of variations (age effect) and hence degree of freedom (Guillemain et al., 2014). In all species except Quail, individuals were categorized as adults when they were trapped for the first time beyond the age of 2 years. In Quail, where individuals may breed during their first calendar year (Nadal et al., 2018), individuals were classified as adults when they reached their second calendar year (i.e., on January 1 of the year following their year of birth, Euring age code ≥ 5). For each species, the whole study period was divided into encounter occasions that lasted one year, starting on the first day of the ringing period in year *t* and ending the day before the next ringing period in year t+1. The beginning of each ringing period was adapted for each species according to its biology and the trapping period (see Table 1). For each bird, individual encounter histories were formatted by reporting events in each encounter occasion. For Pochards which were marked with both a metal ring and a nasal saddle, these events and associated codes were the following: live physical encounter (code: 1), visual encounter/sighting (2), dead recovery (3). For the other species, which were only marked with a metal ring only, live physical encounters were coded as “1” and dead recoveries were coded as “2”. For all species, the ringing event was the first event and was coded as “1”. The absence of encounter at a given occasion was coded as “0”. For the sake of simplicity, only one event was reported per occasion in the individual encounter histories. Therefore, for the Pochard for which individual could be physically and visually encountered during the same occasion, only the physical (more reliable) encounter was retained. Finally, all death events in all encounter histories were shifted by one occasion as in (Guillemain et al., 2014).

### Model structure

Multi-event models (Pradel, 2005) implemented in the E-SURGE software (Choquet et al., 2009) were used. Hidden individual heterogeneity in survival was modelled using the approach developed by Guillemain et al. (2014), which assumes that individuals in a population belong to one of two groups: a first group of individuals with a high survival rate, and a second group of individuals with a lower survival rate. Groups also differ in their probability of live encounter, but have similar probabilities of dead recovery (Guillemain et al., 2014). Accordingly, our models were parameterized to assign individuals to one of these two groups: a newly ringed individual had a π probability of being assigned to the high survival group and a 1-π probability of being assigned to the low survival group. Individuals in the high survival group survived with an annual probability φ, while those in the low survival group survived with a αφ probability. Here, α was a scale coefficient (0 ≤ α ≤ 1) accounting for the difference in survival rates between the two groups. When π = 1 or when α = 1 the population is considered as strictly homogeneous.

The other parameters in our models included the probability of being physically encountered alive in a given year (noted *p*), for the Pochard only the probability of being visually encountered alive in a given year (noted *v*) and the probability of being recovered and reported as dead (*d*) (see Appendix A for a visual representation of the various parameters used in our models).

A set of three matrices was built to describe these multi-event models: a first matrix describing the assignment of new individuals to one of the two survival groups, a transition matrix describing the survival of individuals from one occasion to the next, and an event matrix depicting the detection probabilities (Figure 1). The matrices used in E-SURGE were the same as those shown in Figure 1, except for the transition matrix which was divided in two because the software does not allow the multiplication of two terms within the same matrix (see Appendix A).

**Figure 1.**
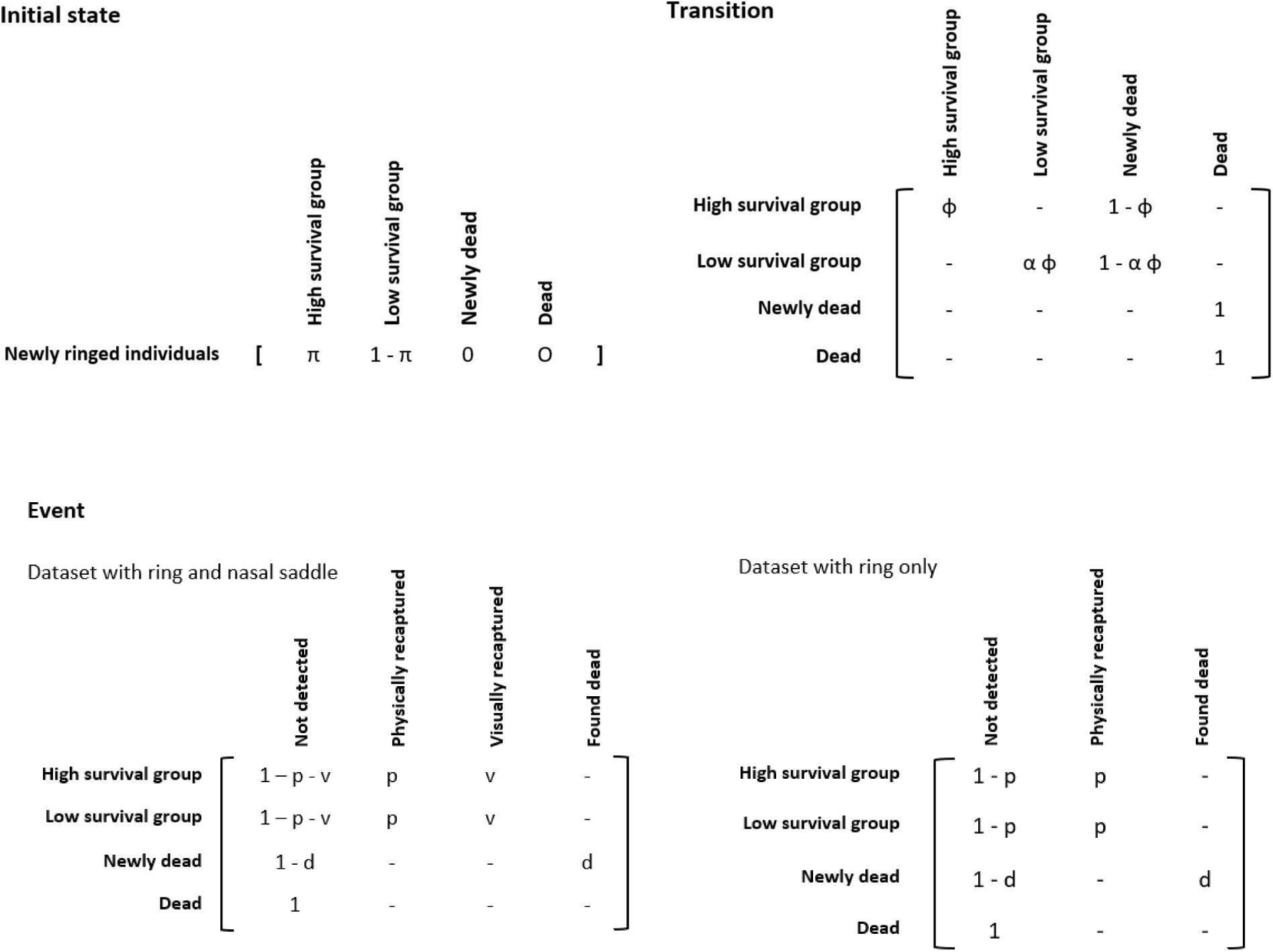
Matrices describing the multi-event model used in our study. π: probability that a newly ringed individual is assigned to the high survival group (1-π for the low survival group); φ: annual survival rate; α: scale coefficient (0 ≤ α ≤ 1) accounting for the difference in survival rates between the two groups; *p*: live physical encounter (recapture) probability; *v*: live visual encounter probability (Pochard only, which were fitted a nasal saddle in addition to the metal ring); *d*: ring recovery probability (conditional on death).

### Initial models and GOF testing

Before conducting the analyses, we assess whether the structure of initial models adequately fit our datasets. Indeed, initial models should include a parameter structure that reflects the main sources of variation of the parameters. In the present study, our initial models accounted for differences among occasions (*time effect*) and between sexes (*sex effect*). However, preliminary fits of complex models, which simultaneously incorporated these effects, led to estimation failures due to data scarcity. Therefore, models were simplified so that their initial structure allowed the various parameters to be properly estimated. In the first step, the complexity of the effects imposed on detection parameters was reduced by assuming no difference between sexes. Unfortunately, even this procedure did not fully resolve the problems of estimating parameters related to survival (π, ɸ and α) when initial models incorporated variation among occasions. Consequently, initial models were simplified by considering that these parameters differed between sexes but were constant over time. For most datasets, the initial model, hence, took the form π(s) ɸ(s) α(s) *p*(t) *v*(t) *d*(t) (see Table 2 for the notation of the effects), except for modern Teal and for Quail. For modern Teal, the model had to be simplified further by keeping the dead recovery probability constant over time (i.e., π(s) ɸ(s) α(s) *p*(t) *v*(t) *d*(.)). For Quail, initial models lacked differences between sexes as our dataset only included males owing to the capture method using calls during the breeding season (i.e., π(.) ɸ(.) α(.) *p*(t) *v*(t) *d*(t)).

**Table 2 -.** Effects tested on the various demographic parameters during the modelling process.

|  |  |
|---|---|
| X (.) | The parameter was constant |
| X (s) | The parameter was subject to a sex effect (two factors: female and male) |
| X (t) | The parameter was subject to a time effect (differences between years as categories) |
| X (tlin) | The parameter was subject to a time effect under a linear trend |
| X (s + t) | The parameter was subject to a sex effect in addition to a time effect |

Goodness of fit (GOF) testing of the initial models was performed using U-Care (Choquet & Pradel, 2003). Full details about this GOF testing approach and the corresponding results are provided in Appendix A. When evidence of a lack of fit was found, a variance inflation factor was calculated, and both standard errors of estimates and AICc values (i.e., QAICc) were adjusted accordingly.

### Model selection and estimates

AICc criteria were used to compare sets of competing models with alternative structures to initial models. We started with a global and fully parametrized model, compatible with the biology of the studied species, and then decreased its parametrization towards a more parsimonious model (using a step-down approach, (Lebreton et al. 1992)). When two models had a delta (Q)AICc of less than 2 units, the parsimony method was applied and we selected the less complex model (Lebreton et al., 1992). If two models had a delta (Q)AICc<2 and the same degree of complexity (same number of estimable parameters), the model with the greater weight (Q)AICc was selected. In order to avoid problems of local minima that could have biased model selection, models were run using 3 random initial values and 3 minimum convergence cycles (a convergence cycle was stopped only when 400 iterations provided similar values of deviance, Choquet & Nogué, 2011, section 10.1).

In the first step, alternative parameterizations related to detectability (*p*, *v*, *d*) were investigated, while keeping the structure of the other parameters in their initial form. These alternative structures constrained parameters as either constant among occasion or different between sexes. Because of estimability issues for some parameters (see previous section), a sex effect was sequentially added or a time effect removed for one parameter at once, starting with *d*.

The selected parameterization for *d* was then used to evaluate alternative parameterizations for *v* (in Pochard). Finally, parameterizations for *p* were assessed while keeping previously selected constraints for *d* and *v*. An overall decline in dead bird ring reporting rates over the recent past decades has been evidenced in UK (Robinson et al., 2009), and suspected for Teal in France (Guillemain et al., 2011). For the modern Teal dataset, where estimability issues prevented considering a categorial time effect on *d*, it was therefore investigated whether considering a linear time trend could be a better alternative parameterization (*d*(tlin), Table 2). The relevance of such a time trend was assessed using both AICc criteria and Likelihood-ratio tests (Grosbois et al., 2008, p. 370). In a second step, and once the initial parameterizations for *p*, *v* and *d* was set, we tested for alternativeparameterizations of π (i.e., π(.) *vs*. π(s)). In a last step, we simultaneously tested the parameterization of ɸ and α; indeed, even if the constraints of the software force to separate these parameters into two matrices, they are linked mathematically. The parameterizations tested were: ɸ(s) α(s), ɸ(.) α(s) or ɸ(s) α(.). Parameterization ɸ(.) α(.) was not assessed because it does not test for hidden heterogeneity beyond the sex (i.e., hidden heterogeneity), so that any recorded difference could simply reflect higher survival of one of the two sexes. The final models were used to estimate parameters π, ɸ, α, *p*, *v*, and *d* and their confidence intervals at population levels. While there is no *a priori* assignment of a given individual to the high or low survival group, E-SURGE allowed *a posteriori* retrieving individual probabilities π*i* of being assigned to the high survival group.

### Sources of heterogeneity

We investigated whether individual probabilities π*i* were related to individual features including bodymass measured at ringing (in kg), ringing date (Julian date with 1 the first day of January) and year of ringing (indicator of a cohort effect), plus sex when this was kept in the final model of the procedure detailed above. To deal with the fact that probabilities π*i* were not evenly distributed, the probability of belonging to the high survival group was split into two groups: “high survival” (π*i* > 0.75) *vs*. “low survival” (π*i* < 0.25). Such uneven distribution of the data was due to the fact that for most species most individuals had a very high or very low π*i* value, while very few individuals had an intermediate π*i* value. To assess the link between π*i* and individual variables, generalized linear models using the “stats” package were used in R 4.3.1 software, choosing "family = binomial" as the link function.

### Between-individual variance in survival

At the population scale (for each dataset), heterogeneity in survival rate is generated by both π (probability to belong to the high survival group) and α (scale effect for survival rate between the high and low survival groups). To assess between-individual variance in survival (*Vm*) in each species sample, we used the composite estimator of Péron et al. (2016):

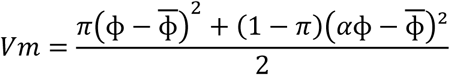

with ɸ being the mean survival rate calculated as ɸ = *π*ɸ + (1 − *π*)*α*ɸ, and π and α the estimates from the best models. We used *Vm* values were used to assess the relationship of *Vm* with the demographic/life-history tactics of the species along the slow to fast continuum. Following Péron et al. (2016), the (log) mean body mass of birds measured at banding was used as a proxy of the position along this continuum. Owing to the modest number of species studied, a Spearman correlation was used to assess the existence of a potential relationship, and phylogenetic inertia was not considered (see also Péron et al., 2016).

## Results

### Goodness of fit

In most cases no lack of fit was detected within the heterogeneous starting model (see Table 3). The only exceptions concerned historical Teal and Blackbird, where the Goodness of fit test turned out to be significant. In both cases, the variance inflation factor was adjusted accordingly (value in Table 3 and details of the test in Appendix A): Blackbird, ĉ = 274/209 = 1.31 and historical Teal ĉ = 537/346 = 1.55.

**Table 3 -.** Goodness of fit test results for each dataset. Estimation of the goodness of fit for the models is done in three steps (see Appendix A), the values shown here are those obtained in the last step. Significant *P*-values at the 5% threshold are in bold.

| **Species and Periods** | **Goodness of fit statistic** | **Degrees of freedom** | ***P*-value** |
|---|---|---|---|
| **Historical Teal** | 537 | 346 | **1.99e-10** |
| **Modern Teal** | 1.3 | 1 | 0.25 |
| **Historical Mallard** | 80 | 89 | 0.73 |
| **Modern Mallard** | 27 | 29 | 0.59 |
| **Pochard** | 15 | 84 | 1 |
| **Snipe** | 126 | 122 | 0.38 |
| **Quail** | 11 | 6 | 0.08 |
| **Blackbird** | 274 | 209 | **0.002** |

### Selected models

The best model retained at the end of the selection process is presented for each species and time in Table 4. The detailed selection process, showing the number of estimated parameters, deviance, AIC, AICc and, if necessary, the weight of the model, are provided in Appendix B. The best model and thus the retained effects on each parameter differed among species, but assignment to the high survival group as well as survival rate of the high survival group were generally subject to a sex effect (in Quail no sex effect was tested because the data set contains only males owing to capture constraints). On the contrary, α and thus the survival of the low survival group was subject to a sex effect in only half of the species and periods. Regarding detection probabilities, dead recovery probability *d* did not vary according to time or sex, with some notable exceptions like the modern Mallard where *d* was much higher in males (0.16) than in females (0.06). For physical and visual encounter probabilities, even though the effects considered could greatly varies between case studies (datasets), a discrete or continuous effect of time in eventually in addition to a sex effect was found (Appendix C).

**Table 4 -.** Best model at the end of the selection process for each species and time period.

| **Species and Period** | **Best model** |
|---|---|
| **Historical Teal** | π(s) ϕ(s) α(.) *p*(t) *d*(s+t) |
| **Modern Teal** | π(s) ϕ(s) α(s) *p*(s+t lin) *d*(.) |
| **Historical Mallard** | π(s) ϕ(s) α(.) *p*(s+t) *d*(.) |
| **Modern Mallard** | π(s) ϕ(s) α(.) *p*(t) *d*(s) |
| **Pochard** | π(.) ϕ(.) α(s) *p*(.) *v*(s) *d*(t) |
| **Snipe** | π(s) ϕ(.) α(s) *p*(s) *d*(s) |
| **Quail** | π(.) ϕ(.) α(.) *p*(.) *d*(.) |
| **Blackbird** | π(.) ϕ(s) α(.) *p*(t) *d*(.) |

Out of the 8 analyzed datasets, 6 yielded α values different from 1 for at least one sex, ranging from 0.93 (0.80-098) in Teal during the historical period to 0.13 (0.03-0.39) in Quail (Table 5). Thus 6 datasets (species or species/periods combinations) out of 8 displayed survival hidden heterogeneity, even if the π values were sometimes high meaning that one of the survival rate categories contained most individuals. Of the 5 species out of 6, displayed survival hidden heterogeneity, the only exception being Blackbird (Table 5). When a difference existed between sexes was evidenced, females generally displayed lower levels of hidden survival individual heterogeneity (differences between the high and low survival groups) than males. In Snipe the between-sex difference in hidden heterogeneity was extreme with females displaying no heterogeneity at all while the α value was 0.81 (0.40 – 0.96) in males.

**Table 5 -.** Survival parameters **ɸ**, probability of assignment to the high survival group **π** survival differential between the high and low survival group **α** and survival of the low survival group **αxɸ** for each species with 95%confidence interval in parentheses.

| **Species and Period** | **Π** | **ϕ** | **α** | **α ϕ** |
|---|---|---|---|---|
| **Historical Teal females** | 0.93 (0.87-0.97) | 0.58 (0.56-0.61) | 0.93 (0.80-0.98) | 0.54 |
| **Historical Teal males** | 0.87 (0.79-0.92) | 0.64 (0.62-0.66) |  | 0.59 |
| **Modern Teal females** | 0.82 (0.65-0.92) | 0.78 (0.43-0.94) | 1.00 (1.00-1.00) | 0.78 |
| **Modern Teal males** | 0.29 (0.16-0.47) | 0.62 (0.44-0.77) | 1.00 (1.00-1.00) | 0.62 |
| **Historical Mallard females** | 0.72 (0.36-0.92) | 0.63 (0.55-0.70) | 0.70 (0.44-0.87) | 0.44 |
| **Historical Mallard males** | 0.97 (0.94-0.99) | 0.58 (0.54-0.62) |  | 0.41 |
| **Modern Mallard females** | 0.81 (0.56-0.93) | 0.66 (0.43-0.83) | 0.64 (0.29-0.89) | 0.43 |
| **Modern Mallard males** | 0.95 (0.85-0.98) | 0.89 (0.50-0.99) |  | 0.57 |
| **Pochard females** | 0.85 (0.72-0.93) | 0.68 (0.64-0.72) | 0.90 (0.36-0.99) | 0.61 |
| **Pochard males** |  |  | 0.77 (0.48-0.93) | 0.53 |
| **Snipe females** | 0.19 (0.10-0.33) | 0.63 (0.58-0.67) | 1.00 (0.00-1.00) | 0.63 |
| **Snipe males** | 0.72 (0.13-0.98) |  | 0.81 (0.40-0.96) | 0.51 |
| **Blackbird females** | 0.10 (0.07-0.13) | 0.46 (0.43-0.50) | 1.00 (0.00-1.00) | 0.46 |
| **Blackbird males** |  | 0.53 (0.49-0.56) |  | 0.53 |
| **Quail (males)** | 0.09 (0.00-0.89) | 0.44 (0.03-0.95) | 0.13 (0.03-0.39) | 0.06 |

Except for Quail whose proportion of individuals assigned to the high survival group (π) was very low 0.09 (although with a wide confidence interval: 0.00 – 0.89), other species had the high survival group (i.e. α ≠ 1) containing the majority of individuals (π values were between 0.67 and 0.97).

### Sources of heterogeneity based on assignation

The analyses yielded conflicting results concerning possible underlying the source of survival hidden heterogeneity (Table 6). For example, the relationship between individual probability of assignment to the high survival group (πi) and body mass was significant for Teal and Mallard of the historical period with Teals with a lower body mass having a higher probability of belonging to the high survival group, whereas the reverse was true for Mallards. In historical Mallard and in Blackbird, individuals ringed towards the end of the season had a higher probability of belonging to the high survival group than conspecifics ringed at the beginning of the season. Historical Mallards individuals belonging to different years of ringing were assigned to different survival groups, although the effect was weaker than for body mass and ringing date. In this case, the individuals ringed during the most recent years had a higher probability of assignment to the high survival group. In the other species (Teal and Mallard of the modern period, Pochard, Quail and Snipe), neither year of ringing nor body mass or date of ringing had a significant effect on the probability of assignment to the high survival group.

**Table 6 -.** Results of binomial models assessing the probability of assignment to the high survival group as a function of individual ringing day (Julian date), body mass, and ringing year. When a sex effect on π was in the best model (see Table 4), this factor was also considered here. Significant effects at *P*<0.05 are indicated in bold.

| **Historical Teal** |  |  |  |  |
|---|---|---|---|---|
| **Coefficients** | **Estimate** | **SE** | **z-value** | ***P*-value** |
| Intercept | -8.919 | 25.346 | -0.352 | 0.725 |
| Ringing day | -0.036 | 0.097 | -0.372 | 0.710 |
| **Sex (M)** | **-0.982** | **0.217** | **-4.519** | **< 0.0001** |
| **Body mass** | **-0.428** | **0.092** | **-4.528** | **< 0.0001** |
| Ringing year | 0.007 | 0.013 | 0.558 | 0.577 |
| **Modern Teal** |  |  |  |  |
| **Coefficients** | **Estimate** | **SE** | **z-value** | ***P*-value** |
| Intercept | -65.709 | 310.399 | -0.212 | 0.832 |
| Ringing day | 0.141 | 0.281 | 0.503 | 0.615 |
| **Sex (M)** | **-6.098** | **0.848** | **-7.192** | **< 0.0001** |
| Body mass | 0.392 | 0.363 | 1.079 | 0.280 |
| Ringing year | 0.035 | 0.155 | 0.225 | 0.822 |
| **Historical Mallard** |  |  |  |  |
| **Coefficients** | **Estimate** | **SE** | **z-value** | ***P*-value** |
| **Intercept** | **109.016** | **38.358** | **2.842** | **0.004** |
| **Ringing day** | **0.532** | **0.186** | **2.867** | **0.004** |
| Sex (M) | 0.476 | 0.371 | 1.282 | 0.200 |
| **Body mass** | **0.582** | **0.151** | **3.867** | **0.0001** |
| **Ringing year** | **-0.058** | **0.020** | **-2.956** | **0.003** |
| **Modern Mallard** |  |  |  |  |
| **Coefficients** | **Estimate** | **SE** | **z-value** | ***P*-value** |
| Intercept | 178.241 | 237.746 | 0.750 | 0.453 |
| Ringing day | -0.154 | 0.403 | -0.382 | 0.702 |
| Sex (M) | 0.873 | 0.592 | 1.475 | 0.140 |
| Body mass | 0.020 | 0.342 | 0.057 | 0.954 |
| Ringing year | -0.087 | 0.118 | -0.735 | 0.463 |
| **Pochard** |  |  |  |  |
| **Coefficients** | **Estimate** | **SE** | **z-value** | ***P*-value** |
| Intercept | 183.369 | 110.208 | 1.664 | 0.096 |
| Ringing day | 0.553 | 0.327 | 1.690 | 0.091 |
| Body mass | -0.334 | 0.271 | -1.236 | 0.216 |
| Ringing year | -0.089 | 0.055 | -1.629 | 0.103 |
| **Snipe** |  |  |  |  |
| **Coefficients** | **Estimate** | **SE** | **z-value** | ***P*-value** |
| Intercept | 9.925 | 38.753 | 0.256 | 0.798 |
| Ringing day | 0.165 | 0.094 | 1.745 | 0.081 |
| **Sex (M)** | **2.160** | **0.252** | **8.568** | **< 0.0001** |
| Body mass | 0.003 | 0.094 | 0.033 | 0.973 |
| Ringing year | -0.007 | 0.019 | -0.340 | 0.734 |
| **Blackbird** |  |  |  |  |
| **Coefficients** | **Estimate** | **SE** | **z-value** | ***P*-value** |
| Intercept | 9.551 | 15.722 | 0.607 | 0.544 |
| **Ringing day** | **0.546** | **0.062** | **8.780** | **< 0.0001** |
| Body mass | 0.051 | 0.031 | 1.629 | 0.103 |
| Ringing year | -0.007 | 0.008 | -0.892 | 0.373 |
| **Quail** |  |  |  |  |
| **Coefficients** | **Estimate** | **SE** | **z-value** | ***P*-value** |
| Intercept | 241.390 | 476.364 | 0.507 | 0.612 |
| Ringing day | 0.145 | 0.736 | 0.197 | 0.844 |
| Body mass | -0.006 | 0.723 | -0.009 | 0.993 |
| Ringing year | -0.124 | 0.236 | -0.525 | 0.600 |

### Interindividual variance in survival (*Vm*)

Figure 2 shows the interindividual variance in survival rate in males of the various species. *Vm* did not decrease with body mass (*rs*=0.19; df=6; *P*=0.66), even though the species with the lowest body mass (Quail) displayed the greatest variance. Species such as Blackbird or Teal had a lower variance than Pochard or Mallard even though their mean body mass is much lower. Concerning females (Figure 3), a trend was marginally significant although opposite to the results of Péron et al. (2016). In fact, in our data set limited to harvested species the variance in hidden survival seemed to increase with body mass (*rs*=0.75; df=5; *P*=0.07), while the opposite was previously recorded for protected species.

**Figure 2:**
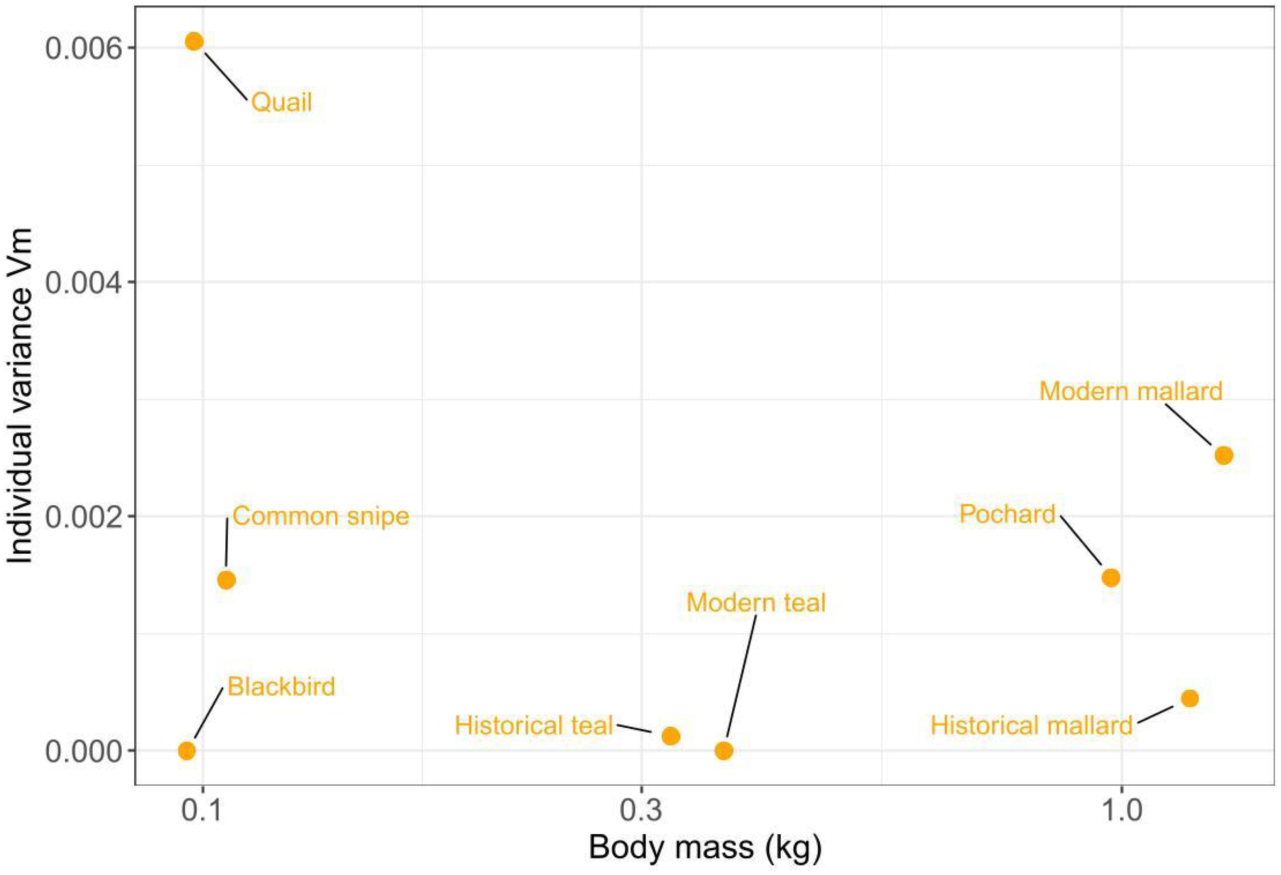
Individual variance for survival rate (*Vm*) as a function of body mass (kg) for male birds. See text for statistics.

**Figure 3:**
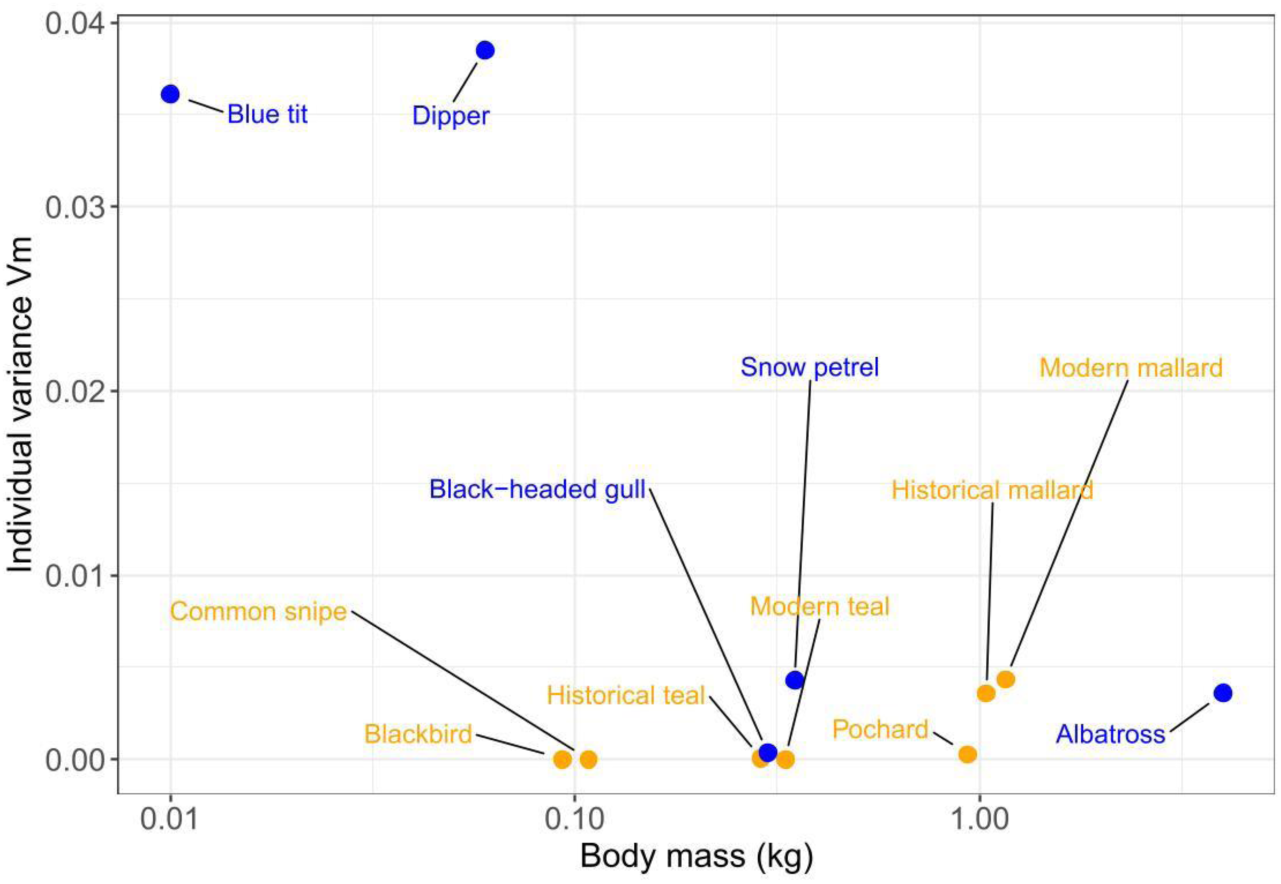
Individual variance for survival rate (*Vm*) as a function of body mass (Kg) in female birds. Game bird species are indicated by orange dots, protected bird species (data from Péron et al. 2016) are indicated by blue dots. See text for statistics.

## Discussion

In most of the hunted species investigated (5 out of 6), we detected two groups of survival heterogeneity. Earlier studies did assess hidden heterogeneity in birds, but most of these studies considered a single species: Black-legged Kittiwake *Rissa tridactyla,* Aubry et al. (2011); Wandering albatross *Diomedea exulans*, Barbraud et al. (2013); Black Brant *Branta bernicla,* Lindberg et al. (2013) Common Teal, Guillemain et al. (2014); Common eider *Somateria mollissima,* Guéry et al. (2017); and, Southern fulmar *Fulmarus glacialoides,* Jenouvrier et al. (2018). Only Péron et al. (2016) and Arnold (2021) simultaneously assessed hidden heterogeneity in large samples (respectively 3 ungulates, 5 protected birds, and 12 North American harvested birds). Both gathered evidence of hidden survival heterogeneity (bimodal variation in survival), although in 6 cases out of 12 only concerning Arnold (2021). Our study covers a large specter of harvested European species, with body mass ranging from ca. 0.1 to 1.3 kg and contrasted ecologies (from purely terrestrial to aquatic species), and migration strategies (spanning from facultative to long-distance migrants e.g., del Hoyo et al., 1992 1994, 1996, 2005). Thus, taken together, studies performed so far suggest that hidden survival heterogeneity is pervasive. Interestingly, many harvested species display hidden heterogeneity although the underlying cause of this heterogeneity is unclear. In our case, the presence of such levels of hidden heterogeneity not necessarily expected because having focused on adults, we could expect that harvesting -either through the non-random removal of vulnerable (frail/low surviving) individuals or by preventing good quality (high surviving) individuals to fully express their potential-had leveled off between-group differences in survival rates. This was clearly not the case, which is consistent with a recent study showing that harvesting as it is practiced on the species investigated here is mostly non-selective (Grzegorczyk et al., 2022).

### Survival estimates

Although our modeling exercise did not intend to provide unbiased survival estimates, most of our survival estimates were within the range of expected values. For adult Teal, our survival estimates –well above 0.50– more or less match those of Guillemain et al. (2014). However, there was a trend towards an increase in female survival over the monitoring period, and the formerly recorded hidden heterogeneity entirely vanished in the modern dataset which yielded very (unusually) high estimates for this species (0.78). Earlier studies have shown that climate change and habitat management have made wintering conditions more favorable, especially recently. Indeed, fewer cold spell were recorded and as result food availability probably improved leading to increasing body condition (Guillemain et al., 2010), which in turn may have being favoring survival.

In Mallard, the heterogeneity increased from the historical to the modern period but was accompanied by an increase in survival in both the low- and high-survival groups. As for Teal, more benign weather conditions accompanied by improved feeding conditions on the wintering grounds (Guillemain et al., 2010), could explain such improvement in survival. Furthermore, Mallards have been experiencing changes in migratory behavior - experiencing short-stopping (sensu Elmberg et al., 2014) and becoming more sedentary (Guillemain et al., 2015)-that could account for the observed increases in survival. Finally, Mallards may have experienced increasing level of hidden heterogeneity owing increases of the practice consisting in the release of farmed individual since about de 1970’s too (Champagnon, 2011). Indeed, although the majority of released individuals experience premature death, some survive until the breeding season (in the Camargue wetlands 34% of the individuals were of captive origin, Champagnon et al., 2016).

Concerning Pochards, the average estimate of survival (0.67) is within the range of the variation observed in previous studies (range 0.61-0.81) including those performed on the same dataset (Blums et al., 1996, Gourlay-Larour et al., 2014, Folliot et al., 2020). Only males of the low survival group did display survival probabilities laying outside of this range (0.53). In the present study, however, we did not detect any between-sex difference in survival in the high survival group, although they seem pervasive in this species (Folliot et al., 2020). Moreover, male of the low survival group did display lower survival than females of the same group although lower female survival is the norm (Folliot et al., 2020).

Our survival estimates of Snipes lay in the upper range of values estimated elsewhere except for males of the low survival group which represent the minority of males (Péron et al., 2013, Petersen et al., 2023). It should also be noted that Snipes display quite high survival probability despite being rather small bodied.

Survival estimates of both Blackbirds and Quails are scarce. Available data suggest that our estimates for Blackbirds would lay in the lower range of the species, that is around 0.50 against more than 0.60 (in Robinson et al., 2010, Robinson et al., 2012). Concerning Quails, our survival estimates (which suffer from too large confidence intervals), are either above previously estimated values for the high survival group (0.44 against 0.27/0.31 in Puigcerver et al., 1992) or well below them, for the low survival, although much less frequent group (0.09). These two species display survival probabilities that are compatible with expectation according to their body mass.

### Inter-individual variance in survival probability and the canalization/buffering hypothesis

Although our study demonstrates that hidden survival heterogeneity is pervasive, the inter-individual variance of survival probabilities, *Vm* (which account for both, the survival differential between the two groups and the proportion of individuals in each group) remains quite low, especially in females. Thus, females which display less variance in than males would be more inclined to buffer survival from environmental stochasticity (Stearns & Kawecki, 1994, Hilde et al., 2020). We hypothesize that such between sex-differences buffering intensity could –at least for long lived species– be explained by higher fitness benefits of demographic buffering for females, whose fitness is highly dependent of future reproductive prospect, than for males (that can sire several females at a time), whose fitness depend more on current than future reproduction. Such a hypothesis is currently difficult to assess largely because many studies focus only on females because they are considered as the limiting sex (in demographic terms). Promoting population dynamics studies in both males and females and assessing inter-individual variability in each sex in a wider range of species and situations, would therefore be relevant, both, to better understand the dynamics of such populations, and to better understand the underlying causes of canalization/demographic buffering. It should be noted that in some species displaying sex-related differences in migratory strategies (differential and partial migration) such as the Pochard, females which migrate further South should be more exposed to harvesting (Carbone & Owen, 1995), and hence should express wider *Vm* than males, which was not observed here.

The probability of assignment to the high survival group (π) was very polarized, with most species showing most high survivors (π>0.72) and a few cases, such as Blackbird and Quail where the high survival group was in the minority (π≤0.10). Arnold (2021), also reported that most species with hidden heterogeneity had a small part of their individuals (3-15%) belonging to low survival group, which displayed very low survival and was termed as a “doomed surplus”. In our study, no such doomed surplus could be evidenced, as the differential in survival between the high- and low-survival group was quite low (range 0.64-0.93), except for Quails (0.13).

According to the canalization/demographic buffering hypothesis, species would tend to buffer (from environmental stochasticity) demographic parameters to which their demography/fitness depends on the most (adult survival for slow species and reproductive success for fast ones, see e.g., Stearns & Kawecki, 1994, Hilde et al., 2020). Contrary, to Péron et al. (2026), our data do not support the buffering hypothesis, because we found a positive relationship between body mass (a proxy of demographic pace) and the variance in survival (*Vm*), where the reverse relationship was expected (Péron et al., 2016). However, our dataset included harvested species only, whereas Péron’s et al. dataset included 3 exploited ungulates and 5 protected birds. Thus, we can hypothesize that harvesting could be the cause of the deviation from the expectation. If our hypothesis if true, therefore, it would mean that the impact of harvesting increases with survival (or body mass/demographic slowness). Our study, therefore, would demonstrate that, as expected, the sustainability of harvesting decreases with life expectancy (e.g., Niel & Lebreton, 2005, Dillingham & Fletcher, 2008, Cooch et al., 2014, Koons et al., 2014).

Finally, despite we used species taken from various bird families with different ecologies, one cannot rule out that the species in our panel were too close in terms of demographic paces to allow a proper testing of the canalization/buffering hypothesis. Indeed, Péron’s et al. (2016) study, included species with body mass ranging from 0.01 to 4 kg in birds and up to reach 170 kg when including ungulates, whereas body mass of species included in our study varied between ca 0.1 and 1.1 only.

### Possible causes underlying (hidden) heterogeneity

Our unexpected results concerning the absence of evidence of demographic buffering (see above) raise the question of the underlying causes of hidden survival heterogeneity. Individual demographic heterogeneity can schematically results from intrinsic factors (e.g., inherent differences of quality/frailty or of demographic strategies/paces of life), or from extrinsic factors such as environmental spatial heterogeneity, and eventually, from purely random demographic processes (Vindenes et al., 2008, Réale et al., 2010, Cooch et al., 2014, Cam et al., 2016, Gimenez et al., 2018). We cannot rule out that part of the heterogeneity observed in our study results, for example, from intrinsic differences between individuals (e.g., poorer body condition, Greenwood et al., 1986; (e.g., poorer body condition, Greenwood et al., 1986, Reinecke & Shaiffer, 1988, Heitmeyer et al., 1993). However, the fact that our findings violate the expectations of the buffering hypothesis begs for the intervention of external causes, in this case, most probably harvesting. Thus, if our hypothesis is true, part of the observed hidden survival heterogeneity would result from (fine grained) spatial differences in harvesting pressure that cannot be *a priori* (explicitly) handled (see Vindenes et al., 2008). However, we must recognize that the heterogeneity was not truly hidden for the Mallard in which individuals could be assigned to a survival group depending on their time of exposure to hunting during the year of ringing, a pattern already stressed by Tavecchia et al. (2001).

While we concur that further investigations are badly needed to better assess heterogeneity, some of the present results (including the aspects above discussed) shed light onto the phenomenon. In Pochards, for example, the probability of physically recapturing the individuals (*p*) was much higher for the low survival group (0.17 against 0.02). This pattern was even more pronounced concerning visual encounters of nasal saddled individuals (respectively 0.57 for females and 0.78 for males vs. 0.11 for females and 0.14 for males). In this species, therefore, the individuals from the high survival group might be either shyer (and hence less exposed to mortality) than the individuals from the low survival group (see Ioannou et al., 2008, Madden & Whiteside, 2014, Miyamoto & Araki, 2020), or would select sites with lower human occurrence (and hence lower hunting pressure). One way to explore this issue would be to correlate personality data (collected before or after releasing the birds) with fitness traits (Madden & Whiteside, 2014).

Another potential extrinsic hidden cause of differential survival, specific to ducks, would be linked to ingestion of lead shot pellets, whose potential to affect survival has been proved (Tavecchia et al., 2001, Guillemain et al., 2007). Indeed, although the use of lead pellet cartridges in wetlands has been banned in many European countries, the problem of lead ingestion seems still acute (Green & Pain, 2016, Béchet et al., 2025).

## Conclusions and perspectives

Overall, the literature is replete of examples showing that hidden heterogeneity is common, if not pervasive in birds (see references above). Here we show that the same conclusion holds for harvested species. However, our results clearly violates the assumption of the buffering hypothesis that predicts decreasing variance in survival with demographic slowness (Péron et al., 2016). Instead, we detected a reverse trend, which in our opinion gives credit to the “exploitability hypothesis”, which predicts increasing impact of harvesting with demographic slowness (e.g., Niel & Lebreton, 2005, Dillingham & Fletcher, 2008, Cooch et al., 2014, Koons et al., 2014). This is an important issue that could be addressed thanks to the implementation of adaptive harvest which is becoming a reality for some species in Europe (Cruz-Flores et al., 2024). Indeed, by playing on harvest rates, we should be able to put to the test the exploitability hypothesis. In the meantime, we badly need implementing studies explicitly accounting for intrinsic causes of heterogeneity such as personality types (Réale et al., 2010), as well as on other extrinsic causes such as lead pellets ingestion or fine-grained variations in predation (hunting) pressure (see above). All of this should be done keeping in mind key theoretical frameworks such as frailty, pace-of-life and exploitability hypotheses (see above). Finally, for statistical purpose we considered two survival groups as many others did before us (see above), while survival may rather be distributed along a continuum as predicted by theory (Dammhahn et al., 2018). As such, however, the approach used in our study and in previous ones, proved very powerful for detecting survival heterogeneity and could prove very efficient e.g., to assess sex-related differences in survival in monomorphic species. Coupled with genetic sexing of a subsample of individuals one would be able to efficiently estimate survival rates for each sex separately. The same approach would also allow assessing the survival consequences of different migratory strategies coexisting in the same population (Gourlay-Larour et al., 2014).

## Acknowledgements

We would like to thank the hundreds of bird ringers and the thousands of field observers and hunters who helped build the birds’ datasets used here. Géraldine Simon is thanked for computerizing the historical dataset. Betty Plaquin, Alexandre Villers, Damien Coreau and Maxime Passerault are thanked for running the modern dabbling duck, blackbird, quail and snipe ringing groups.

